# Thermal plasticity in protective wing pigmentation is modulated by genotype and food availability in an insect model of seasonal polyphenism

**DOI:** 10.1101/2024.02.26.582119

**Authors:** E van Bergen, G Atencio, M Saastamoinen, P Beldade

## Abstract

Developmental plasticity refers to the phenomenon whereby an organism’s phenotype depends on the environmental conditions experienced during development. This plasticity can match phenotype to ecological conditions and help organisms to cope with environmental heterogeneity, including differences between alternating seasons. Experimental studies of developmental plasticity often focus on the impact of individual environmental cues and do not take explicit account of genetic variation. In contrast, natural environments are complex, comprising multiple variables with combined effects that are poorly understood and may vary among genotypes. We investigated the effects of multifactorial environments on the development of the seasonally plastic eyespots of *Bicyclus anynana* butterflies. Eyespot size is known to depend on developmental temperature and to be involved in alternative seasonal strategies for predator avoidance. In nature, both temperature and food availability undergo seasonal fluctuations. However, our understanding of whether thermal plasticity in eyespot size varies in relation to food availability and across genotypes remains limited. To address this, we investigated the combined effects of temperature (T; two levels: 20°C and 27°C) and food availability (N; two levels: control and limited) during development. We examined their impact on wing and eyespot size in adult males and females from multiple genotypes (G; 28 families). We found evidence of thermal and nutritional plasticity, and of temperature-by-nutrition interactions (significant TxN) on the size of eyespots in both sexes. Food limitation resulted in relatively smaller eyespots and tempered the effects of temperature. Additionally, we found differences among families for thermal plasticity (significant GxT effects), but not for nutritional plasticity (non-significant GxN effects) nor for the combined effects of temperature and food limitation (non-significant GxTxN effects). Our results reveal the context dependence of thermal plasticity, with the slope of thermal reaction norms varying across genotypes and across nutritional environments. We discuss these results in light of the ecological significance of pigmentation and the value of considering thermal plasticity in studies of the biological impact of climate change.

## INTRODUCTION

Phenotypic diversity, both within and across species, is shaped by interactions between organisms and their environment, which occur across levels of organization and time scales. On the one hand, environmental conditions play a crucial role in shaping population-level phenotypic frequencies across generations, through the process of natural selection. On the other hand, environmental conditions influence organismal phenotype expression, through the phenomenon of phenotypic plasticity. This environmental dependence of phenotype expression is represented graphically by reaction norms (Schlichting & Pigliucci, 1998), which display phenotype as a function of environmental conditions. Developmental plasticity, in particular, refers to the phenomenon by which the phenotype is influenced by the environmental conditions during development (Beldade et al., 2011; West-Eberhard, 2003). When these conditions anticipate the ecological conditions organisms will face later in life, developmental plasticity can match phenotype to future environment and, as such, help populations cope with environmental heterogeneity (Ghalambor et al., 2007). Such adaptive developmental plasticity is well illustrated by the phenomenon of seasonal plasticity, common in insects (Brakefield et al., 1996; Brakefield & Zwaan, 2011; Simpson et al., 2011), where seasonal conditions lead to the production of alternative phenotypes and life strategies (i.e. seasonal polyphenism). Moreover, it has been argued that developmental plasticity can impact not only the immediate survival but also future adaptation of populations facing environmental perturbation (Bonamour et al., 2019; Reed et al., 2011) or colonizing novel environments (Bilandžija et al., 2020). These ideas underscore the importance of integrating developmental plasticity into research about the biological consequences of climate change (e.g. Barnes, 2021; Rodrigues & Beldade, 2020; Sgrò et al., 2016).

The effects of external environmental conditions on developmental outcomes have been extensively documented across phenotypes and species. Developmental plasticity, i.e. differences attributed to the environment (E), and genotype-by-environment interactions (GxE), i.e. genetic variation for plasticity, are commonly observed, including for butterflies (e.g. life-history traits in Verspagen et al., 2020). While evolutionary genetics distinguishes between genetic effects (G) that are additive versus non-additive (i.e. GxG interactions comprising dominance and epistasis) and make distinct contributions to evolution, the environment is often regarded as an irreducible entity. This perspective overlooks potential environment-by-environment interactions (ExE) that can influence phenotype expression and organismal fitness. Many experimental studies of plasticity focus on analysing single environmental factors that are kept constant throughout an organisms’ lifetime. This approach starkly contrasts with natural scenarios, where environments are multifactorial, including various cues that may act additively on phenotype expression, but also redundantly, synergistically, or antagonistically (Piggott et al., 2015; Westneat et al., 2019). Furthermore, these interaction effects may vary across traits, sexes, and genotypes. Quantifying the effects of multifactorial environments on developmental plasticity is crucial for our understanding of how organisms adapt to changing environments. However, only a limited number of studies have examined ExE and GxExE in insects to assess how reaction norms differ between genotypes and between environments (Bubliy et al., 2013; Guilhot et al., 2020; Kutz et al., 2019; Stoehr & Wojan, 2016; Verspagen et al., 2020). Across seasons, for example, various biotic and abiotic factors change more or less in concert, including temperature and food quantity/quality. Both of these are key properties of the developmental environment and can affect various traits (e.g. Clissold & Simpson, 2015; Lazarević et al., 2023), but the impact of their interaction on developmental outcome is poorly characterised.

We propose to examine the effects of multifactorial environments using a model system of adaptive thermal plasticity. Temperature exhibits spatial variation (e.g. altitudinal and latitudinal gradients), as well as temporal fluctuations (e.g., circadian and seasonal changes). Its influence on plasticity and adaptation has been well-documented, especially for ectotherms (Colinet et al., 2015). *Bicyclus anynana* butterflies have become a valuable experimental model of the ecological significance and physiological basis of thermal seasonal plasticity (Brakefield et al., 2009). The temperature experienced during pre-adult stages determines the production of distinct phenotypes associated with alternative seasonal strategies for survival and reproduction. In its natural habitat in Eastern and Southern Africa, alternating wet- and dry-seasons differ in various environmental factors, including temperature and food plant availability (see scheme in van Bergen et al., 2016). Butterflies from one versus the other season differ in wing pigmentation patterns and in a series of life history and behavioural traits that make up a so-called plasticity syndrome (Brakefield & Zwaan, 2011). Seasonal variation in eyespot size (Brakefield and Reitsma, 1991; Brakefield et al., 2009) and colour of eyespot rings (van Bergen & Beldade, 2019) underlie seasonal differences in the strategy to avoid predators. While the larger and conspicuous marginal eyespots of wet-season butterflies deflect predator attention away from the more vulnerable body, the duller colour patterns of dry-season butterflies provide camouflage against the background of dry leaves. In the laboratory, dry and wet season phenotypes can be produced by rearing individuals at lower versus higher temperatures, respectively. While food quantity has been shown to also influence some of the life history traits that underlie *B. anynana* seasonal differences in pace of life (Saastamoinen et al., 2013), there is currently no information regarding the effects of food quantity and its interaction with temperature on the wing patterns involved in alternative seasonal strategies for predator avoidance.

In this study, we quantified the effects of the environment (E), genotype-by-environment interactions (GxE), environment-by-environment interactions (ExE), and genotype-by-environment-by-environment interactions (GxExE) on the size of the seasonally plastic eyespots of *B. anynana*. Specifically, we focused on investigating the combined influence of temperature (T) and food availability (N) on eyespot size in multiple genetically distinct families. We measured wing size and eyespot size in female and male progeny from 28 father-mother pairs whose eggs were split into four treatments (Fig 1A and 1B) corresponding to the full factorial combination of variation in temperature (20°C versus 27°C, used to induce development of dry-versus wet-season phenotypes, respectively) and variation in food level (control versus food limitation, another ecological factor likely to change in anticipation of the onset of the dry-season). We found evidence for thermal plasticity as well as nutritional plasticity, and for temperature-by-nutrition interactions (TxN) on eyespot size. Food limitation resulted in relatively smaller eyespots and tempered the effects of temperature on eyespot size. Additionally, we found differences among families for thermal plasticity (significant GxT effects), but not for nutritional plasticity nor for the combined effects of temperature and food quantity (non-significant GxN and GxTxN effects, respectively). We discuss these results in terms of the importance of considering multifactorial environments and genetic variation for environmental sensitivity in studies of how organisms respond to environmental change.

**Figure 1:**
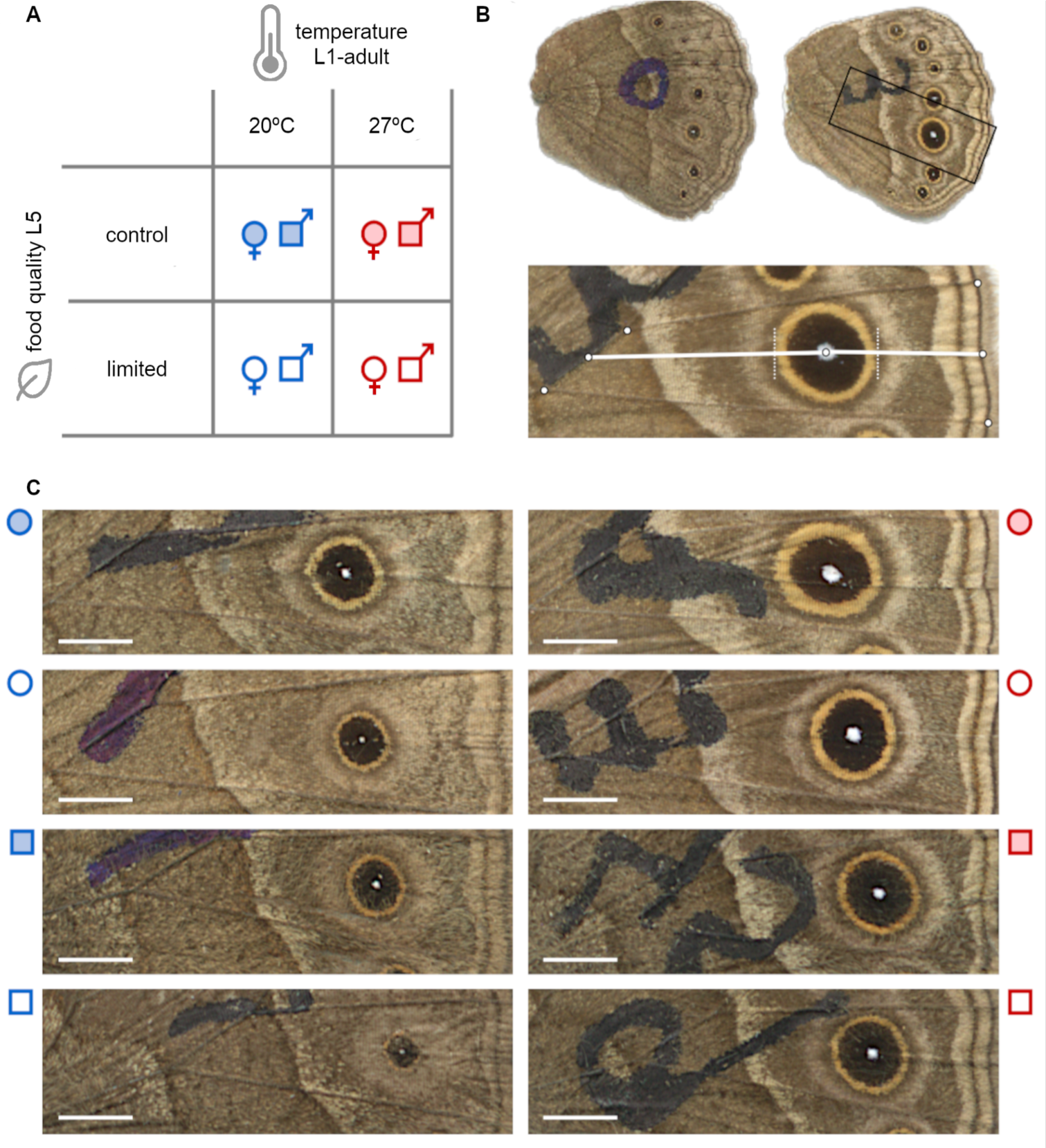
Experimental design and target traits. **A**) Treatments: Progeny from 28 couples from the outbred stock of B. anynana were split into four developmental conditions, corresponding to the combination of two temperatures (20°C vs 27°C, used to induce development of dry- vs wet-season morphs, respectively) and two levels of food availability (control vs limitation, with the latter mimicking what is the scenario for the dry-season). **B**) Phenotyping: Images of the ventral surface of the hindwing of two representative females from extreme conditions: development at 20°C with food limitation (left) versus development at 27°C with ad libitum food. Black marks on wings were added with permanent marker pen as part of the processing of butterflies for analysis of life-history traits in a previous study (Saastamoinen et al., 2013). Insert below wings shows the transect (white line) defined by landmarks (white circles; see Materials and Methods) on the wing compartment featuring the hindwing eyespot typically used in studies of eyespot size plasticity in B. anynana. The length of the transect was used as a proxy of wing size, and the eyespot diameter along the transect (portion delimited by perpendicular dotted lines) as a proxy for eyespot size (cf. van Bergen and Beldade, 2019). **C**) Examples of target eyespots in adult females and males from the distinct developmental conditions (legend next to each image cf. panel A). White size bar corresponds to 2 mm.

## RESULTS

We measured wing and eyespot size, two traits known to differ between the wet- and dry-season morphs of *B. anynana*, in progeny from 28 families reared under four conditions. These conditions represented the full-factorial combination of two temperatures (20°C and 27°C) and two levels of food access (control and limited) (Figure 1). Differences in phenotype between same-family individuals developed at 20°C vs. 27°C reflect thermal plasticity, and this thermal plasticity can be calculated for control and for limited food access conditions. Conversely, differences in phenotype between same-family individuals developed under control vs. limited food access correspond to nutritional plasticity, and this can be calculated for individuals developed at 20°C and for individuals developed at 27°C. As such, for each family, we calculated four reaction norms for each of the sexes.

Our dataset allowed us to address several open questions about plasticity in *B. anynana* eyespot size, a trait whose seasonal polyphenism is relatively well understood in terms of both its ecological significance and physiological underpinnings. Is there plasticity in relation to food access? Are the combined effects of temperature and food access additive or non-additive (Figure 2)? Is there genetic variation for the effects of temperature, food availability, and their interaction (Figure 3)? What is the relationship between environmental sensitivity under different conditions (e.g. thermal plasticity under control versus limited food access), to different cues (temperature or food quantity), and across the sexes (Figure 4)?

**Figure 2:**
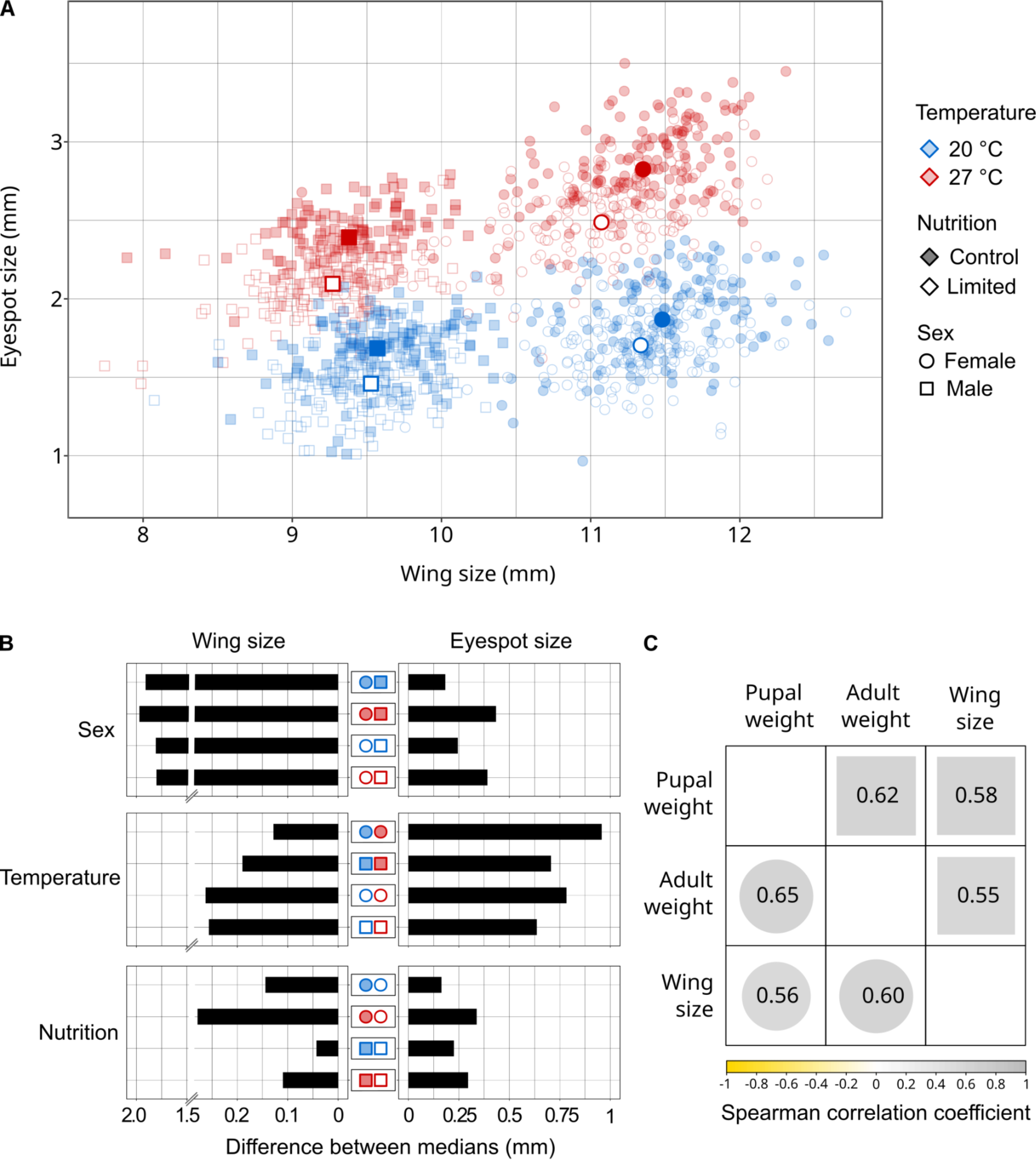
Variation in eyespot size and wing size. **A)** Values of eyespot and wing size for all individuals measured (smaller symbols), and corresponding median values per sex and treatment (larger symbols). We tested for phenotypic differences between sexes and across treatments, where treatments refer to the four possible combinations of two temperature and two food access levels. For wing size, we found significant differences between sexes (F_1,1157_=5347.74, p<2e-16) and between treatments (F_3,1157_=36.83, p<2e-16) but no significant treatment-by-sex interaction (F_3,1157_=0.8584, p=0.462). For eyespot size, using wing size as covariate, we found significant differences between sexes (F_1,1156_=4.0905, p=0.043) and between treatments (F_3,1156_=1626.45, p<2e-16), as well as a significant treatment-by-sex interaction (F_3,1156_=22.9041, p=2.1e-14). **B)** Summary of effects of treatments and sexes: absolute value of difference in median values for wing (left) and eyespot (right) size between pairs of groups (center; legend cf. panel A). **C)** Spearman correlations between individual measures of body size, including transect length as our proxy of wing size (Figure 1B) and pupal and adult weight characterized in a previous study (Saastamoinen et al., 2013). Analysis was done separately for females (circles on bottom half) and males (squares on top half). All correlation coefficients displayed are significantly different from zero (adjusted p-value<0.05).

**Figure 3:**
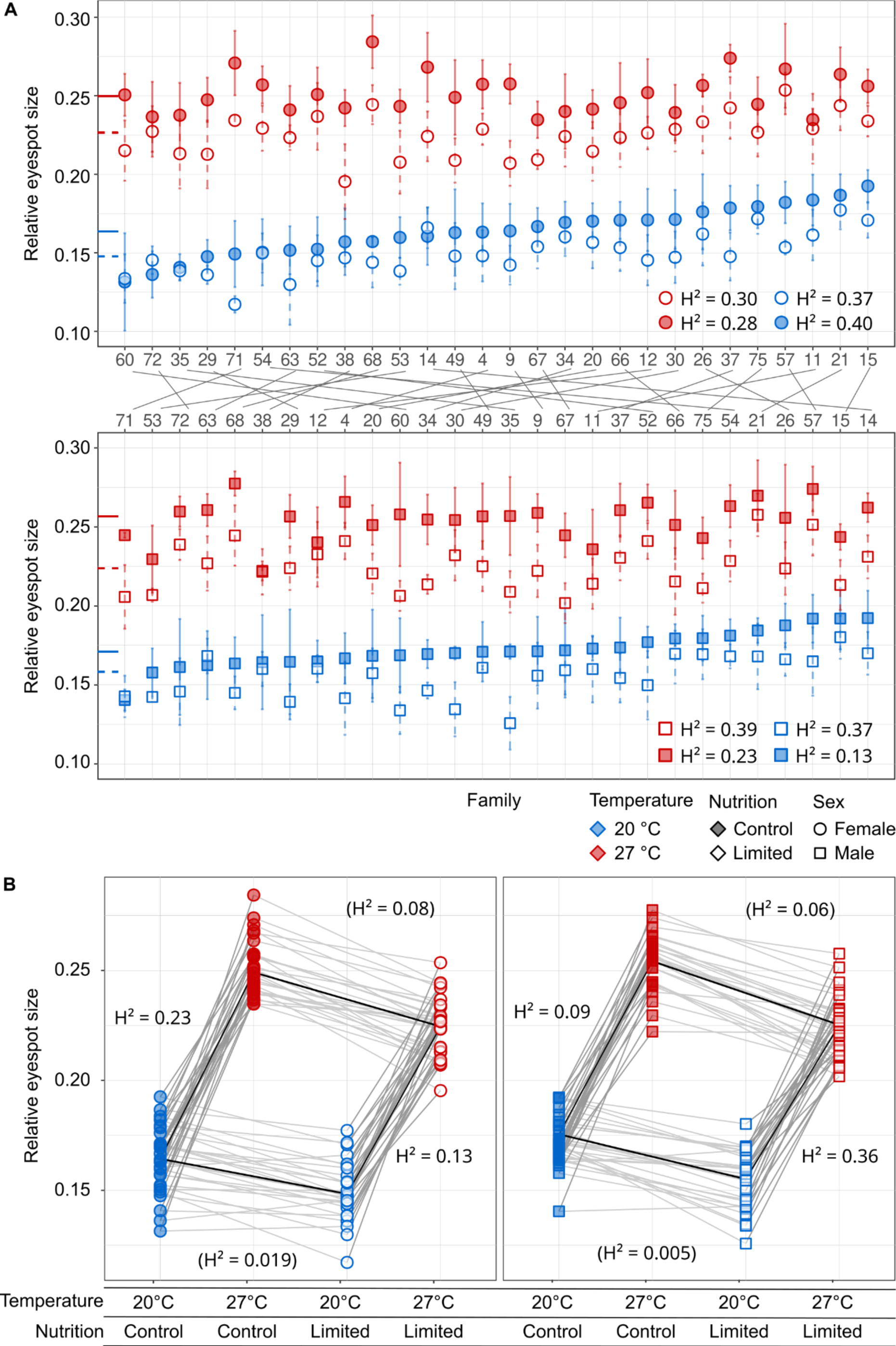
Genetic variation for thermal and nutritional plasticity in eyespot size. **A)** Relative eyespot size for females (top, circles) and males (bottom, squares), with symbols representing median values (with standard deviations) for the different families. Along the X-axis families appear in ascending order of median relative eyespot size for individuals reared at 20°C with ad libitum food (filled blue symbols). Estimates of broad-sense heritability (H^2^) for relative eyespot size under different environmental conditions are included in each plot. Lines between top and bottom plots connect the same families, illustrating sexual dimorphism in relative eyespot size and in plasticity therein. **B)** Reaction norms across temperatures (symbols of different colours) and across food availability conditions (filled versus empty symbols). Symbols represent median relative eyespot size for each of the different families, thinner lines connect family medians across environments and represent the reaction norms for each family, thicker lines connect median values across all families and represent median reaction norms. The slopes of the regression lines corresponding to the reaction norms are shown in Figure 4A. Estimates of broad-sense heritability for plasticity are inserted next to the corresponding reaction norms, and shown inside brackets for nutritional reaction norms, which were not significantly different between families (non-significant GxN in Table 1).

**Figure 4:**
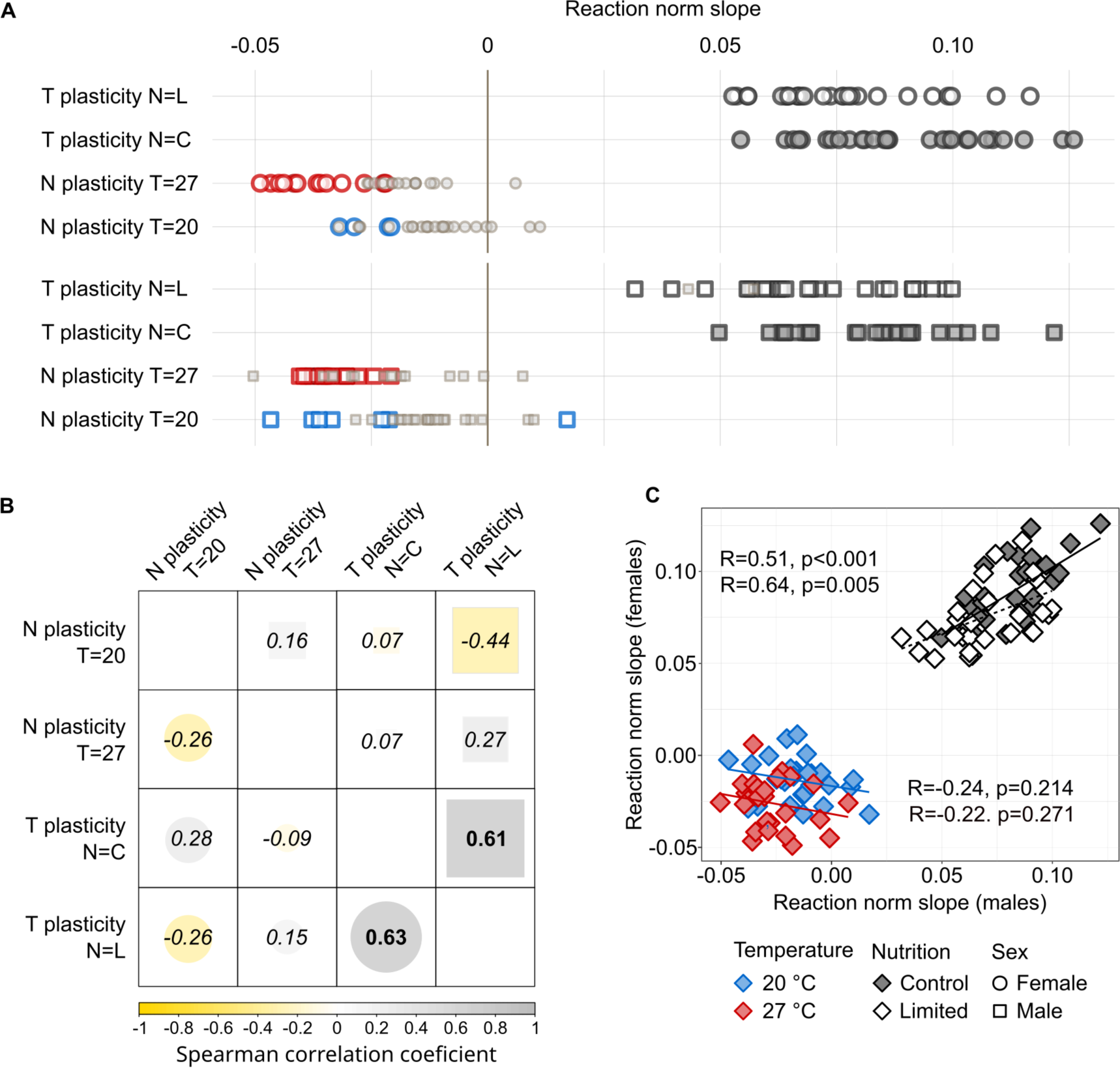
Correlation between different types of plasticity across families. **A)** Slope of reaction norms for each family (symbol) and sex (females as circles, males as squares): thermal plasticity for the two food conditions and nutritional plasticity for the two temperature conditions. Smaller symbols in grey represent slopes that were not statistically different from zero (adjusted p-value>0.05), corresponding to families that were not plastic. Slope and corresponding adjusted p-values in table S2. **B)** Spearman correlations between slopes of reaction norms: nutritional (N) and thermal (T) plasticity for the different temperature (T=20°C versus T=27°C) and food quantity (N=Control versus N=Limited food access) conditions, respectively. Correlations were calculated separately for females (circles, below the diagonal) and males (squares, above the diagonal). Symbol colour reflects correlation coefficient cf. colour gradient underneath correlogram. Numbers correspond to the actual correlation coefficients; in italics when not significantly different from zero (adjusted p>0.05), and bold when statistically significantly different from zero (adjusted p<0.05). **C)** Relationship between female and male reaction norm slopes. Symbols represent the slope of reaction norms for each family; thermal reaction norms as grey symbols (filled for control food access, empty for limited food access) and nutritional reaction norms in colour (blue for 20°C, red for 27°C). Lines represent linear regression and adjacent text represents Spearman correlation coefficient (R) and corresponding p-value (p).

### Nutritional plasticity and temperature-by-nutrition “epistasis”

Both wing size and eyespot size differed between sexes and between treatments (Figures 1C and 2A). Wings and eyespots were smaller in males relative to females, and in individuals exposed to food limitation relative to those with continued *ad libitum* food access. In terms of temperature effects, as had been described in previous studies of plasticity in *B. anynana* pigmentation (reviews in Beldade & Monteiro, 2021; Beldade & Peralta, 2017; Brakefield et al., 2009), individuals reared at warmer temperature had smaller wings and larger eyespots than those reared at cooler temperature.

Our analysis included transect length as a proxy for wing size (Figures 1B). This measurement was positively correlated with other proxies of body size quantified in a previous study of the same target individuals (Saastamoinen et al., 2013) (Figure 2C). As documented for various insects (e.g. Adamo et al., 2012; Davidowitz et al., 2003; Karl et al., 2011; Lee & Roh, 2010), adults developing at higher temperatures or experiencing food limitation were smaller than those from lower temperatures or with *ad libitum* food access, respectively. The temperature effects on body size are consistent with the populational patterns described by Bergmann’s rule (see Shelomi, 2012), which states that organisms tend to be larger where climate is colder (namely, higher altitudes and latitudes). In our dataset, the effect of food availability on wing size was greater for females than for males, and resulted in reduced sexual dimorphism under food limitation relative to control nutritional conditions (Figure 2B).

We then focused on analysing differences in eyespot size relative to wing size across temperature (T) and food (N) conditions, and across families (G), separately for males and females. For both sexes, we found significant differences between developmental temperatures (T) and food availability conditions (N), as well as significant temperature-by-nutrition (TxN) interactions (Table 1). While thermal plasticity in relative eyespot size in *B. anynana* has been amply documented (reviews in Beldade and Monteiro, 2021; Beldade and Peralta, 2017), the effect of food quantity on eyespot size was never described. We found that food availability, which was shown to affect some but not all life history traits involved in alternative seasonal pace-of-life modes (Saastamoinen et al., 2013), affected eyespot size, a trait involved in alternative seasonal strategies for predator avoidance (see van Bergen & Beldade, 2019). The significant TxN interactions we documented correspond to non-additive effects between temperature and food availability. Nutritional plasticity was reduced when individuals were reared at cooler temperature, and, conversely, thermal plasticity was reduced when caterpillars experienced food limitation (Figure 2B).

**Table 1:**
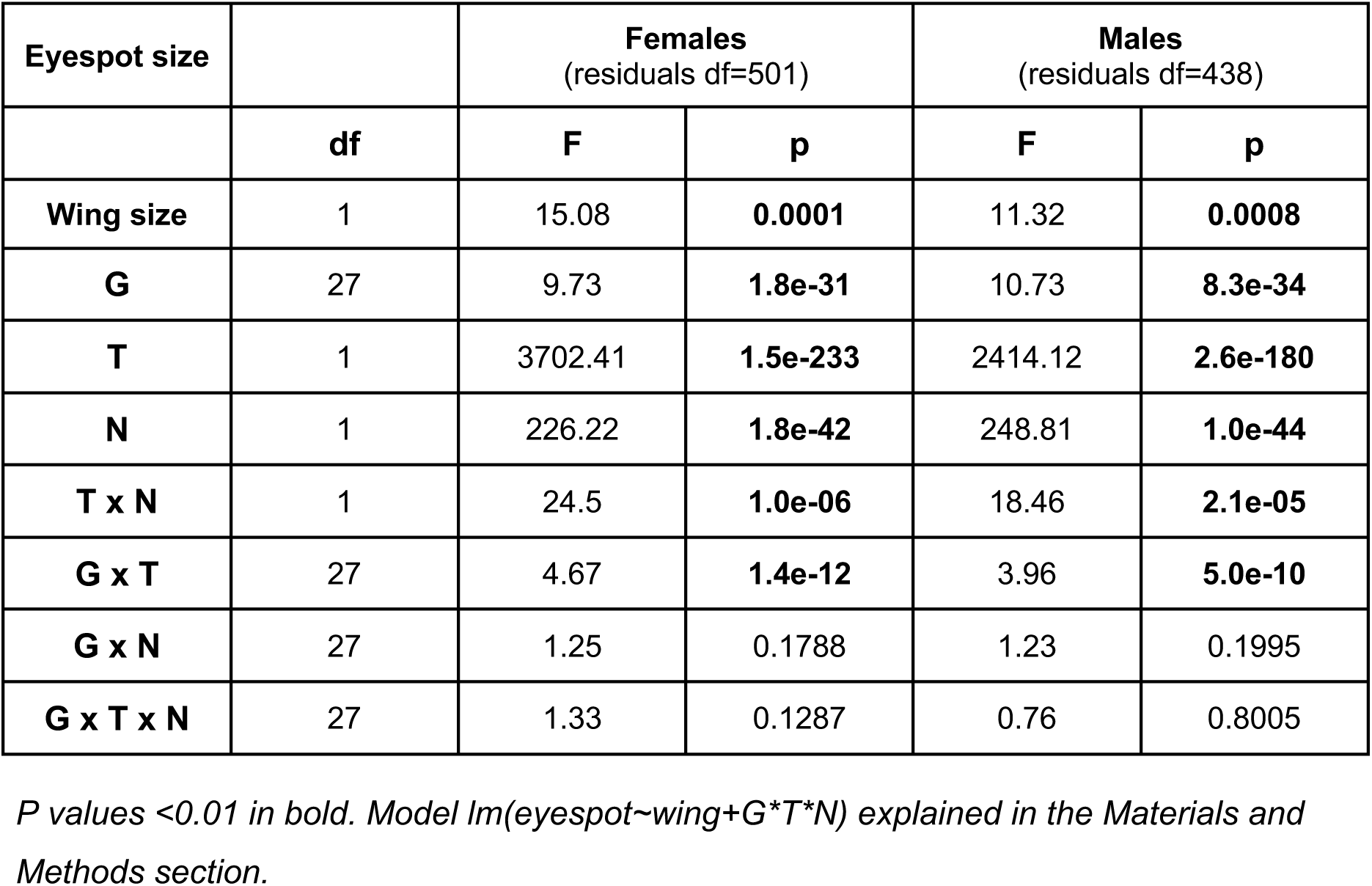
Statistical analysis of variation in eyespot size in relation to genotype (G), temperature (T), and food availability (N)

### Genetic variation for eyespot plasticity

We observed differences among families for relative eyespot size (significant G effects in Table 1), with broad-sense heritability of 20-40%, except for males from cooler temperatures and limited food conditions (Figure 3A). Additionally, we found significant differences between families for thermal plasticity (GxT), but not for nutritional plasticity (GxN) nor for TxN interactions (GxTxN) (Table 1). This corresponds to genetic variation for the slope of thermal reaction norms, but not for the slope of nutritional reaction norms nor for the response to temperature-by-nutrition effects (Figure 3B). Consistently, broad-sense heritability estimates were higher for GxT (varying between 9% and 36%) than for GxN (varying between 1% and 8%) (Figure 3B). Across the 28 families and for both sexes, thermal reaction norms were steeper than nutritional reaction norms (Figure 3B, Figure 4A). Consistently, the slopes of thermal reaction norms were generally significantly different from zero, while the slopes of nutritional reaction norms were often not (Figure 4A, file S2).

To explore to what extent plasticity is a general property of a genotype, versus one that differs between sexes and between environmental variables, we investigated the correlation between the slopes of the different reaction norms calculated for each of the families (Figure 4). For both males and females, we found a significant positive correlation between the slopes of thermal reaction norms across food availability conditions (Figure 4B): i.e. families with steeper thermal reaction norms under *ad libitum* food tended to also have steeper thermal reaction norms under food limitation conditions (Figure 4A). We also found significant positive correlations for the slope of thermal reaction norms between males and females: i.e. families for which thermal reaction norms for females were steeper were also those with steeper reaction norms for males (Figure 4C). On the other hand, and consistent with nutritional reactions norms not differing significantly across families (Table 1), we found no significant correlation between slopes of nutritional reaction norms across temperatures, nor between the slopes of thermal versus nutritional reaction norms (Figure 4B), and no significant correlation between slopes of nutritional reaction for females versus males (Figure 4C).

## DISCUSSION

Colour and colour patterns are key aspects of the interaction between organisms and their environment. On the one hand, pigmentation plays important roles in how organisms deal with biotic and abiotic variables, including roles in thermoregulation, UV protection, sexual signalling, defence against parasites, and avoidance of predators (e.g. Bybee et al., 2012; Hanley et al., 2013; Olofsson et al., 2010). On the other hand, pigmentation development is affected by environmental variables, including host plant (Boyle & Start, 2020) and ambient temperature (e.g. Brakefield et al., 2009; Lafuente et al., 2021, 2023; Mateus & Beldade, 2022). We focused on the characteristically thermally plastic eyespots of *B. anynana* butterflies (reviews in Beldade & Monteiro, 2021; Beldade & Peralta, 2017; Brakefield et al., 2009). Eyespot size and colour differ between the wet- and dry-season morphs in association with their distinct strategies for predator avoidance (see van Bergen & Beldade, 2019). However, while both temperature and food availability vary seasonally, only the effect of temperature on eyespot patterns had been documented previously. Here, we investigated the combined effects of temperature and food access on *B. anynana* eyespot size in progeny from distinct families to test for the existence of nutritional plasticity and for the genetic and environmental dependence of thermal plasticity.

### Food availability varies across seasons and affects pigmentation development

The natural habitat of *Bicyclus* butterflies is characterized by alternating wet and dry seasons, which differ in a variety of ways. The difference in vegetation cover is believed to be a key ecological feature behind the distinct seasonal strategies for predator avoidance (Brakefield & Reitsma, 1991). While butterflies from the dry season have inconspicuous wing pigmentation patterns and avoid predator attention through camouflage against the background of dry foliage, butterflies from the wet season have larger eyespots that deflect predator attack towards the wing margin and away from the vulnerable body. A decrease in ambient temperature anticipates the arrival of the dry season and induces the development of the dry-season morph. Conversely, an increase in ambient temperature anticipates the arrival of the wet season and induces the development of the wet-season morph. As such, temperature works as an inductive cue (cf. Lafuente & Beldade, 2019) that predicts future vegetation cover, and induces the development of adult phenotypes better adjusted to it.

We posited that food availability also varies seasonally, with food shortage becoming more common with the decrease in available green vegetation at the onset of the dry season. As such, food plant availability could be a reliable predictor of vegetation cover and selection could have favoured developmental plasticity in relation to food availability. Our results showed a significant effect of food access on eyespot size (Table 1), with food limitation during larval life inducing the development of smaller eyespots (Figures 1C and 2A), which is the typical phenotype of the dry season morph. This is consistent with a previous study demonstrating the effect of food plant quality on *B. anynana* eyespot size (Kooi et al., 1996; Singh et al., 2020), and supports the idea that nutritional plasticity for eyespot size is adaptive in relation to seasonal variation. On the other hand, we found that food limitation resulted in smaller bodies, which is the opposite of what is characteristic of the dry season morph. As such, rather than being adaptive in relation to the specific conditions of each season, nutritional plasticity in body size is advantageous by enabling the production of a functional adult in spite of food shortage. Previous studies had documented the effect of food availability on other seasonally variable life history traits in *B. anynana* (Saastamoinen et al., 2013), as well as various life history traits in other insect species (e.g. Choy et al., 2023; Kutz et al., 2019), but not on pigmentation patterns.

While both decreasing temperature and food shortage anticipate the onset of the dry season, it is difficult to assess to what extent one cue might be a more reliable predictor than the other (see also discussion in Halali et al., 2021). Food limitation, while more directly connected to vegetation cover, is also more spatially heterogeneous (see Arnoldini et al., 2012). On the other hand, the information value of temperature depends on climate types and cues other than temperature could replace it on induction of alternative seasonal morphs (Roskam & Brakefield, 1999). Importantly, it is difficult to assess to what extent the food quantity treatments we used experimentally are realistic representations of natural scenarios. There are various reasons for this difficulty. First, while our larvae endured food stress for a single period during the last larval instar, in nature it is possible to have multiple food stress events across instars, interrupted by periods of feeding. Second, for both experimental and natural settings, it is unclear whether and how much larvae can compensate food shortage by overeating when food becomes available again. In this experiment, for example, the larvae that experienced food limitation took longer to complete development, but still did not reach the same body size as their control counterparts (Saastamoinen et al., 2013). Third, in natural situations both the quantity and quality of food likely vary more or less in concert, with availability of fewer and lower quality food plants in the dry season. Finally, it is worth mentioning that while we used a full factorial design in our experiment (Figure 1A), in nature the different combinations of temperature and food quantity might not be equally likely. In fact, we expect the onset of the dry season to be characterized by lower temperature and food shortage (akin to our treatment with limited food and 20°C), and the wet season to be characterized by higher temperature and *ad libitum* food (akin to our treatment with control food conditions and 27°C). These were, in fact, the experimental conditions that resulted in smallest and largest eyespots, respectively (Figure 1C, 2), consistent with what is expected to be favoured in each of the corresponding natural seasons. It has been argued that plasticity in relation to multiple environmental cues could be beneficial either because plastic responses might work better in synergy or because they complement each other under different circumstances (Nielsen & Papaj, 2022).

### Thermal plasticity varies with genetic and environmental context

Thermal plasticity in *B. anynana* eyespot size was relatively well characterized in terms of its physiological basis and its ecological significance. In contrast, we knew relatively little about genetic variation for this thermal plasticity and even less about its dependence on food availability. Our data revealed that thermal plasticity in eyespot size depends on both genetic and environmental context, consistently across sexes.

By comparing reaction norms across 28 families that represent standing genetic variation in our study population (Figure 3A), we found significant GxT interactions (Table 1), which reveal that thermal reaction norms differ between families (Figure 3B). Previous studies had shown variation in thermal reaction norms for *B. anynana* eyespot size across families (Wijngaarden & Brakefield, 2001; Windig, 1994), across strains bearing spontaneous mutations of large effect on pigmentation (Mateus & Beldade, 2022), and across geographical populations (Jong et al., 2010). Here, we analysed standing genetic variation available in a large outbred captive population (Brakefield et al., 2009), presumably the type of genetic variation that could feed evolutionary change in levels of plasticity (see Lafuente & Beldade, 2019). Our results revealed abundant genetic variation for the slope of thermal reaction norms, in contrast with previous claims of limited genetic variation for thermal plasticity in this experimental population of *B. anynana* (Oostra et al., 2018; Wijngaarden & Brakefield, 2001).

By comparing temperature effects (T) under different situations of food access (N), we found significant TxN interactions (Table 1). This corresponds to combined effects of T and N that are not additive, and reveals that the extent of thermal plasticity depends on nutritional context (Figure 2A, 3A). Specifically, we found that thermal plasticity was reduced when caterpillars experienced food limitation (Figure 2B). In the same manner that quantitative genetics uses “epistasis” to describe non-additive interactions between alleles at different loci, these non-additive interactions between distinct environmental variables can be thought of as “environmental epistasis” (see Samir et al., 2015). This type of temperature-by-nutrition interactions had been documented for different life history traits in different species (e.g. (Kingsolver et al., 2006; Lee & Roh, 2010; Stillwell et al., 2007), but, to our knowledge, not for pigmentation development.

### Developmental plasticity and adaptation to seasonal and directional environmental change

The evolution of plasticity, under whatever ecological conditions that might favour or disfavour it, requires the existence of heritable variation for the slopes of reaction norms (e.g. Lafuente & Beldade, 2019). In this study, we found evidence for genetic variation for the slope of thermal reaction norms (GxT effects and estimated of H^2^ up to around 30%), but not for the slope of nutritional reaction norms (no GxN effects and corresponding H^2^ closer to 0%) (Table 1 and Figure 3B). It is unclear to what extent the lack of genetic variation for nutritional plasticity reflects the action of selection, which could have fixed alleles responsible for specific levels of plasticity either as direct target of selection or as a correlated response to selection on other traits. Moreover, we found significant positive correlations between thermal plasticity under control versus limited food conditions (Figure 4B), and between thermal plasticity in males versus females (Figure 4C), suggesting that selection on plasticity in one setting will result in changes in plasticity across settings. On the other hand, we found that our study families did not differ in their response to the combined effects of T and N (no GxTxN). In contrast, segregating genetic variation in combined response to temperature and food quality was documented for *D. melanogaster* life-history traits (Choy et al., 2023).

Seasonal plasticity in *Bicyclus* butterflies is characterised by a suite of traits that change in concert to enable different strategies for survival and reproduction (see van Bergen et al., 2017). These traits include functionally connected pigmentation, life histories, and behavioural traits, whose concerted changes across seasons make up what is referred to as a “seasonal syndrome” (see Brakefield & Zwaan, 2011; Rodrigues et al., 2021). Despite the functional integration of different plastic traits, various previous studies showed that thermal reaction norms can differ between traits and between sexes (Mateus et al., 2014; Muller et al., 2019; Oostra et al., 2011, 2014; Rodrigues et al., 2022; Rodrigues et al., 2021). The previous study that generated the panel of butterflies we analyzed here revealed nutritional plasticity for some but not all life-history traits that contribute to seasonal differences in pace-of life (Saastamoinen et al., 2013). We focused on wing patterns, which are associated with seasonal differences in predator avoidance and documented previously unknown nutritional plasticity and context-dependence of thermal plasticity for eyespot size. Taken together, these results hint at a complex picture of the making of natural phenotypes that must integrate across external environmental inputs and their genetics to produce coherent whole-organism phenotypes (see Forsman, 2015). It has been previously argued that plastic traits might affect each other’s response to environmental cues; either by affecting how the environmental cue is received by the organism or by affecting the response to that cue (Nielsen & Papaj, 2022).

The focus on thermal plasticity is especially pertinent in the context of the biological impact of climate change, which entails global and directional change in temperature averages and temperature extremes. Thermal plasticity can evolve in response to the selection imposed by climate change, but it can also impact both the persistence and future adaptation of populations experiencing it. A number of review and research papers have focused on the importance of including thermal plasticity in studies of climate change (Bonamour et al., 2019; Kaunisto et al., 2016; Oostra et al., 2018; Paull et al., 2015; Reed et al., 2011; Rodrigues & Beldade, 2020; Westneat et al., 2019), including considering effects of multifactorial environments (Rodrigues & Beldade, 2020; Rodrigues et al., 2021). Despite great interest in understanding the effects of multiple stressors in the context of climate change (Piggott et al., 2015), studies investigating such effects remain scarce outside the realms of plants or aquatic systems (Kaunisto et al., 2016). Furthermore, most studies focus solely on environmental conditions that are considered universally and individually stressful, neglecting that what constitutes a stress can vary depending on the genetic and environmental context. This is aligned with what we found: thermal plasticity in eyespot size differed between genotypes and between levels of food availability. The context-dependence of plasticity underscores the value of including more complex experimental scenarios that can better illustrate the breadth of possible natural scenarios. Indeed, theoretical work has shown that analysing plastic responses in isolation might lead to erroneous conclusions relative to what happens in natural populations, which are under the influence of an array of environmental variables (Chevin & Lande, 2015).

## MATERIALS AND METHODS

### Butterflies and experimental conditions

For this study we used the butterfly specimens from a previous full-factorial split-brood experiment (Saastamoinen et al., 2013) examining variation in various life-history traits in relation to two thermal environments, two nutritional treatments, and 28 families (Figure 1A) from an outbred captive population of *B. anynana* (see Brakefield et al., 2009). In short, first instar larvae from each family were split into two rearing temperatures (20°C and 27°C) with *ad libitum* feeding (on young maize plants). On the first day of the fifth and final instar, larvae within each temperature treatment were randomly assigned into one of two nutritional treatments: food limitation and control. Individuals were transferred to Petri dishes with either fresh maize leaves (control) or a piece of agar (to provide humidity) but no maize leaves; henceforth referred to as “food limitation”. These nutritional conditions were kept for 2 days in the warmer conditions and 3 days in the colder conditions, after which *ad libitum* feeding was restored. After eclosion, the adults were sacrificed and their wings were dissected and stored in individual entomological envelopes until phenotyping.

### Phenotyping adult wing size and eyespot size

One ventral hindwing of each individual (N=1165) was scanned using a digital scanner (Epson V700) and the resulting images were analysed with a set of custom-made interactive Mathematica notebooks (see van Bergen and Beldade, 2019). Two contiguous transects were defined by five landmarks on the wing compartment bearing the eyespot that is typically used to characterise seasonal polyphenism in *Bicyclus* and related genera (van Bergen et al., 2017). Total length of the transects was used as a proxy for wing size. Two additional landmarks, specifying the outer limits of the golden eyespot ring along the transect lines, were used to define the eyespot diameter and were used as proxy for eyespot size (Figure 1B).

### Data analyses

All statistical analyses were conducted in R v 4.3.2 (R Core Team, 2018) and RStudio (Posit team, 2023). The *tidyverse* package suite (Wickham et al., 2019) was used for the data preparation, analysis and plotting. The packages *ggplot2* (Wickham, 2009) and *lmtest* (Zeileis & Hothorn, 2002) were used to assess normality and heteroscedasticity, through QQ plots and the Brausch-Pagan test, respectively, before fitting the models for thermal and nutritional response. The package *lme4* (Bates et al., 2015) was used to test for phenotypic differences between treatments by producing linear mixed models. The package *VCA* (Schuetzenmeister & Dufey, 2022) was used to estimate all the variance components. The package *stats* (R Core Team, 2018) was used to calculate the correlations and corresponding adjusted p-values (Holm correction for multiple comparisons). The R code and corresponding results are available as supplementary file S3.

We first analysed all data together and tested for differences in wing and eyespot size between sexes (*sex* as fixed factor with 2 levels) and developmental conditions (*condition* as fixed factor with 4 levels, corresponding to the full-factorial combination of two temperatures and two food access levels). The corresponding R syntax was *lm*(*wing∼condition*sex)* and *lm*(*eyespot∼wing+condition*sex*, with *wing* size as covariate, respectively. We then focused on eyespot size and analysed females and males separately to assess for effects of family (genotype, *G*; fixed factor with 28 levels), temperature (*T*; fixed factor with two levels), and food availability (*N*; fixed factor with two levels), using wing size as covariate. The corresponding R syntax was *lm(eyespot∼wing+G*T*N)*.

Slopes of reaction norms for each family and sex were calculated based on the linear regression fit to eyespot size data on environmental condition. For each family and sex, we obtained slopes of four reaction norms: thermal reaction norms under control food access, thermal reaction norms under limited food access, nutritional reaction norms at 20°C, and nutritional reaction norms at 27°C. Adjusted p-values were calculated with the Holm-Bonferroni adjustment for multiple tests (as implemented in the *stats* package for R (R Core Team, 2018).

We estimated broad-sense heritability (H^2^) for relative eyespot size as *H^2^=σ^2^_A_/(σ^2^_A_ + σ^2^_W_)*, where *σ^2^_A_* is the among-family variance and *σ^2^_W_* is the within-family variance. We estimated broad-sense heritability for relative eyespot size plasticity as proposed in Scheiner and Lyman (1989), *H*^2^=*σ*^2^*_G_*_*E_/*σ*^2^*_TOTAL_*, where *σ*^2^*_G_*_*E_ and *σ*^2^*_TOTAL_* are the variance associated with the *genotype-by-environment* interaction components and the total variance, respectively.

We calculated Spearman correlations between individual values for three different proxies of body size: transect length, our measurement of wing size, and pupal and adult weight from a previous study using the same individuals (Saastamoinen et al., 2013). We also calculated correlations across families between slopes of the four different reaction norms, separately for females and males (Figure 4B), and between the sexes, separately for the four reaction norms (Figure 4C). P-values for correlation coefficients were adjusted for multiple comparisons using Holm correction (as implemented in the *stats* R package; R Core Team, 2018).

## Notes

**Funding:** Portuguese science funding agency, *Fundação para a Ciência e Tecnologia*, research grants PTDC/BIA-EVF/0017/2014 and PTDC/BIA-EVL/0321/2021 to PB.

**Conflict of interests:** None

### Competing Interest Statement

The authors have declared no competing interest.

